# Soft topographical patterns trigger a stiffness-dependent cellular response to contact guidance

**DOI:** 10.1101/2022.01.25.477731

**Authors:** Jordi Comelles, Vanesa Fernández-Majada, Verónica Acevedo, Beatriz Rebollo-Calderon, Elena Martínez

## Abstract

Directional migration is involved in multiple physiological and pathological processes. Among other external signals, the architecture of the extracellular matrix can trigger directed cell migration through a phenomenon known as contact guidance: cells elongate, align, and migrate along the direction set by aligned extracellular matrix fibers. This process involves the orientation of focal adhesions, actin, and tubulin cytoskeleton along the direction of those fibers. Contact guidance has been extensively studied on stiff materials with topographical grooved patterns. However, how it translates to softer physiologically relevant compliances is not known. Here we show that substrate stiffness modulates the cellular response to topographical contact guidance. We found that for fibroblasts, while focal adhesions and actin responded to topography independently of the stiffness, microtubules showed a stiffness-dependent response that regulates contact guidance. On the other hand, both clusters and single breast carcinoma epithelial cells displayed stiffness-dependent contact guidance migration, leading to more directional and efficient migration when increasing substrate stiffness. These results suggest that both matrix stiffening and alignment of extracellular matrix fibers cooperate during directional cell migration, and both should be accounted when studying processes such as cancer cell invasion.

**Teaser:** Changes in the stiffness of topographical patterns modify how mesenchymal and epithelial cells perform contact guidance.

## Introduction

The ability of a cell or group of cells to migrate directionally as a response to external cues is crucial in multiple physiological and pathological processes (*1*–*3*). For example, directional cell migration is needed for neural crest and primordial germ cell migration in development, for immune response, wound healing, or for cell invasion in cancer metastasis among other processes. Mostly, directed cell migration has been attributed to *chemotaxis*, the well-studied ability of cells to follow chemical factors dissolved in the environment. However, directed cell migration can also be driven by mechanical cues (*durotaxis*) (*4*) or even geometrical features. This last form of directed cell migration is known as contact guidance (*5, 6*). Contact guidance results from the presence of micron and sub-micron aligned structures formed by the cellular rearrangement of extracellular matrix protein fibers (typically collagen and fibronectin). Such protein rearrangement can be found *in vivo* in the basal membrane of healthy tissues where it is known to assist tissue formation (*7, 8*). On the other hand, fiber alignment can also be found in the stromal tissue around solid tumors (*9*), where it is identified as a negative prognostic factor (*10*). In this situation, aligned fibers in the extracellular matrix regulate cell extrusion and invasion (*11*). Despite the outcome of contact guidance on cell migration is well known, how the geometrical features are sensed by cells and transduced in directional migration is not fully comprehended.

*In vitro*, contact guidance has been retrieved by culturing cells on grooved substrates. There, cells elongate, align, and migrate in the direction of the grooves mimicking the *in vivo* phenotype on top of collagen fibers. Topographically induced contact guidance has been extensively reported for several cell types: fibroblasts (*12*–*18*), epithelial (*12, 18*–*20*), endothelial (*21, 22*), neuronal (*12, 23, 24*), and muscle cells (*25, 26*), as well as stem cells (*27, 28*) and cancer cells (*29*–*34*). Early on, microtubule and actin cytoskeletons were observed to align with the topographical grooves (*13*–*15, 19, 27, 34*–*36*). Later on, it was reported that grooves lead to the alignment of focal adhesions and the activation of Focal Adhesion Kinase (FAK) and downstream signaling to reorganize actin cytoskeleton (*19*–*21, 31, 33*). Therefore, the cell contractility machinery seems to be playing a part, involving the Ras homolog family member A (RhoA) pathway (*21, 26, 28, 31*). However, contact guidance has been found to be both contractility-dependent (*21, 25, 28, 31*) but also contractility-independent (*22, 37*). Thus, the role of intracellular contractile forces in contact guidance is still controversial and remains unclear. In part, the reason is that most of the reported work so far has been performed onto topographically modified stiff materials such as thermoplastic polymers and polydimethylsiloxane (PDMS) (*19*–*22, 25, 26, 28, 31, 33*). These materials have elastic modulus values of ∼ 10^6^ kPa, which are much harder than the ones encountered in soft tissues *in vivo*, either in physiological or pathological conditions (*38, 39*), which range from 0,1 to 100 kPa. Within this scenario, there is the need to study contact guidance while acknowledging substrate stiffness.

Up to date, contact guidance studies on soft materials have been hampered by the lack of proper microfabrication methods to create well-controlled micron and sub-micron features on such materials. In fact, only a few examples can be found in literature where topographical features are introduced on soft substrates (*34, 37, 40, 41*). These studies have shed some light on the effect induced by topographical grooves on cancer cells, fibroblasts and T-cells in soft environments. Fibroblasts showed elongation and directional migration on aligned soft matrices (*37*), cancer cells responded differently to soft (2.3 kPa) than to stiff (50 kPa) topographical grooves (*34*), and T-cells experienced contact guidance on soft (16 kPa) but not on stiff (50 kPa) substrates (*40*). However, whether a reduction of stiffness would abrogate directional migration through the decrease of cell tractions, or an increase of stiffness would enhance directional response of epithelial cells are questions yet to be solved. Thus, the systematic study of contact guidance within the *in vivo* stiffness range is crucial to explore new strategies to tackle scenarios where topographically induced directed migration takes place *in vivo*. Here, we study the response of contact guidance associated hallmarks, cell elongation, alignment and directional migration, to systematic changes of the substrate stiffness covering physiological, pathological and non-physiological values (Figure. 1a). To that end, micron-size grooves onto polyacrylamide (PAA) hydrogel substrates were produced by using a recently developed microfabrication method (*42*). Our results show that mesenchymal-like cells exhibit stiffness-independent directional migration by contact guidance by modifying the cytoskeleton elements involved in a stiffness dependent manner. On the contrary, single breast carcinoma epithelial cells exhibit stiffness-dependent directional migration by contact guidance, with and increased efficiency upon matrix stiffening. Importantly, not only single cells but also cell clusters are shown to perform directional collective migration upon matrix stiffening. Altogether, these findings highlight the importance of substrate stiffness in contact-guidance associated migration and open new avenues to investigate cancer cell invasion accounting for the cooperation between matrix stiffening and collagen fiber alignment.

**Fig. 1.**
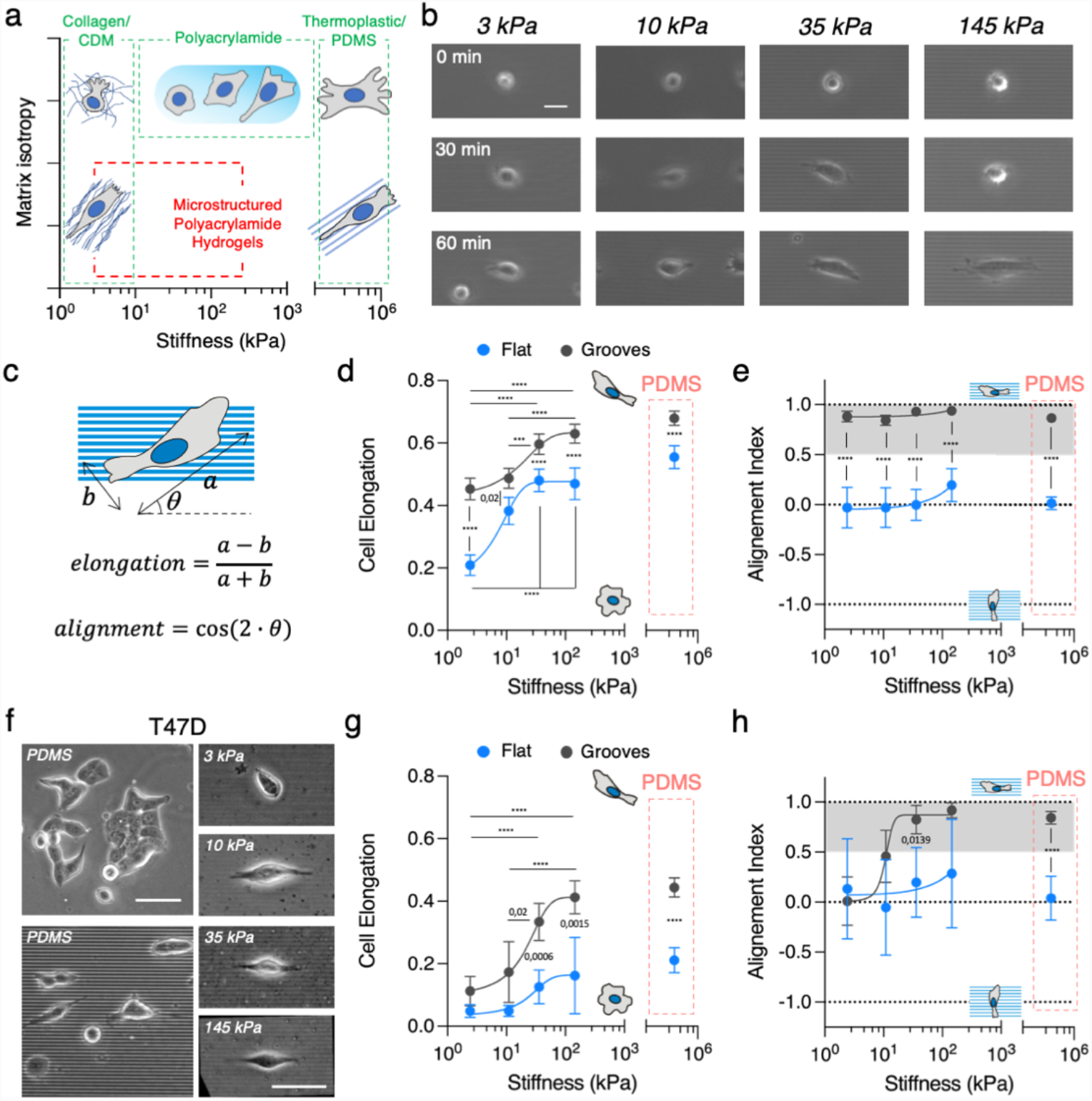
Substrate stiffness modulates topographical contact guidance. (**A**) Schematics of matrix isotropy across a wide range of substrate stiffness. (**B**) Snapshots of NIH 3T3 fibroblasts adhering, elongating and aligning on grooves of different stiffness. Scale bar, 20 μm. (**C**) Scheme depicting cell long axis a, short axis b and the angle between the long axis and the grooves θ. (**D**) Cell elongation and (**E**) cell alignment index as a function of the substrate stiffness. (**F**) T47D cells spread on flat and grooves PDMS substrates. T47D single cells spread on polyacrylamide gels with topographical grooves and ridges of increasing stiffness. Scale bar, 50 μm. (**G**) Cell elongation and (**H**) Alignment index as a function of substrate stiffness. See Tables S1 and S2 for the number of cells and experiments. Data points (Mean ± CI) were fitted as an eye-guide. Statistical significance was assessed by Tukey’s tests (**D, G** and **H**) and Kruskall-Wallis’ test (**E**).

## Results

### Cell elongation and alignment with topography during contact guidance change as a function of substrate rigidity

To explore the impact of substrate stiffness on topographical contact guidance, we fabricated substrates containing 2 μm wide and 1 μm deep topographical grooves and ridges made of PAA gels of different stiffness (3 – 145 kPa) by Capillary Force Lithography (*42*). Because gels swell, the spacing between ridges was slightly smaller for the PAA replicas than for the original PDMS mold used, but overall the microfabrication technique employed was successful in providing well-defined topographies even for the softest gels (Fig. S1a,b). We then coated these grooved substrates and analogous flat, “non-patterned” counterparts, with fibronectin, and we first seeded mesenchymal-like cells, NIH 3T3 fibroblasts, onto them. As soon as cells started spreading on the PAA gels, they elongated and aligned in the direction of the grooves (Fig. 1b and Movie S1), thus showing topographical contact guidance features like those found on standard PDMS grooved substrates (Fig. S1c). Next, we quantitatively characterized contact guidance associated hallmarks on cell morphology, namely cell area, elongation, and alignment with the grooves (see Methods section for details), as a function of substrate rigidity (Fig. 1c). We found cell area increasing with substrate stiffness until reaching a maximum value on PDMS. This trend was observed on the topographically patterned substrates (Fig. S1d) and also on flat counterparts (as expected (*43*)). When quantifying cell elongation, we first observed that NIH 3T3 fibroblasts on flat and soft (3k Pa) PAA gels were roundish, but they broke symmetry and became ‘front-back’ polarized (*44*) when stiffness increased (Fig. 1d). This intrinsic tendency of elongate on flat substrates when stiffness increases was further enhanced by the presence of the grooves, which were also capable to break symmetry and induce elongation onto soft 3 kPa substrates. Finally, while cell population showed no overall alignment (alignment index ∼0) on flat substrates for any of the tested stiffnesses (Fig. 1e,), the cells long axis were strongly aligned with the groove’s direction (alignment index ∼1), for all the grooved substrates tested, independently of their rigidity (Fig. 1e). Overall, mesenchymal-like NIH 3T3 fibroblasts showed topographical contact guidance features on their morphology when cultured on grooves of different stiffness. Remarkably, these could be retrieved even for soft (3 kPa) substrates mimicking the physiological stiffness attributed to the extracellular matrix of soft tissues.

We were then wondering if non-mesenchymal cells such as epithelial cells, which are not polarized ‘front-back’ naturally (*45*), would also undergo topographical contact guidance on soft substrates. To address this question, we employed T47D epithelial cells, a breast carcinoma cell line. These cells are reported to respond to topographical patterns on stiff substrates as a function of their adherens junction (E-cadherin) activity (*33*), but their behavior onto soft patterned substrates is unknown. First, we seeded single T47D cells on stiff substrates (PDMS) and we were able to retrieve cell elongation and alignment guided by the topographical patterns (Fig. 1f). Then, we seeded single T47D cells onto the PAA grooved gels of different stiffness. Interestingly, T47D epithelial cells showed topographical contact guidance features in their morphology (elongation and alignment) that were heavily dependent on the stiffness of the substrate (Fig. 1f). First, despite T47D cells adhered poorly onto soft gels, when adhered, cell elongation was enhanced by the grooves in a stiffness dependent manner (Fig. 1g), breaking their natural circular shape on stiffer gels (35 – 145 kPa). And second, cell alignment with the direction set by the topographical pattern became stiffness dependent as well (Fig. 1h), being less evident on the softest gels and in opposition to what we observed for fibroblasts. Thus, our experiments revealed that epithelial cells do experience topographical contact guidance features on soft substrates but, contrary to what we saw for the mesenchymal-like cells, cell elongation and alignment is strongly modulated by the rigidity of the substrate.

### Cell contractility regulates contact guidance on soft topographical grooves and ridges

To address the role of contractility in topographical contact guidance, we used our experimental set-up to account for the potential effects of substrate stiffness in this phenomenon. First, given that strongly contractile cells such as fibroblasts can make large deformations on soft substrates (*46, 47*), we checked if groove deformation could be observed in our experiments. Indeed, NIH 3T3 fibroblasts were able to significantly deform the topographical patterns on soft PAA gels (3 – 10 kPa), but not on stiffer ones (35 – 145 kPa) (Fig. 2a). To dig further into the molecular actors behind these observation and, since it has been shown that focal adhesions align with topographical patterns on stiff substrates (*19, 33*), we decided to characterize focal adhesion distribution on the different stiffness. To do that, we used fluorescence immunostaining of paxillin and quantified focal adhesion size and alignment with the topographical grooves and ridges. Representative images showed that on grooved substrates fibroblasts developed focal adhesions localized along the direction of the grooves and ridges (Fig. 2b). Regarding the focal adhesion size, mean area increased with substrate stiffness on flat substrates but remained roughly constant on grooved substrates (Fig. 2c). Moreover, focal adhesion sizes were similar for the flat and grooved substrates when they were stiff (145 kPa PAA and PDMS), but they differed when substrates were soft (3 kPa), being significantly higher for the grooved gels. This suggests that grooves enlarged focal adhesions on soft substrates. Concerning focal adhesion alignment, it should be noticed that since fibroblasts were naturally elongated on flat surfaces, alignment was analyzed *vis-à-vis* the main cell axis direction. For the patterned substrates, however, the data plotted were analyzed *vis-à-vis* the groove direction (Fig. S2). On flat surfaces, ∼ 50% of the cell focal adhesions were oriented in the direction of the cell main axis (*θ*_*FA*_ < 30º) for all the substrates (Fig. 2d). As an isotropic, non-aligned distribution would lead to 33% of the focal adhesions with *θ*_*FA*_ < 30º, this demonstrates that the defined main cell axis is indeed a signature of symmetry breaking characteristic of this cell type that favors focal adhesion alignment. On the patterned substrates, ∼ 70% of focal adhesions were oriented in the direction of the grooves for all the substrates, thus demonstrating that focal adhesions alignment does not depend on the stiffness of the substrate. So, despite the deformations of the grooves caused by cell contractility on the softest substrates, the physical restriction imposed by the topographical pattern to focal adhesions was effective in breaking symmetry for all the stiffness range, and led to focal adhesion elongation and alignment.

**Fig. 2.**
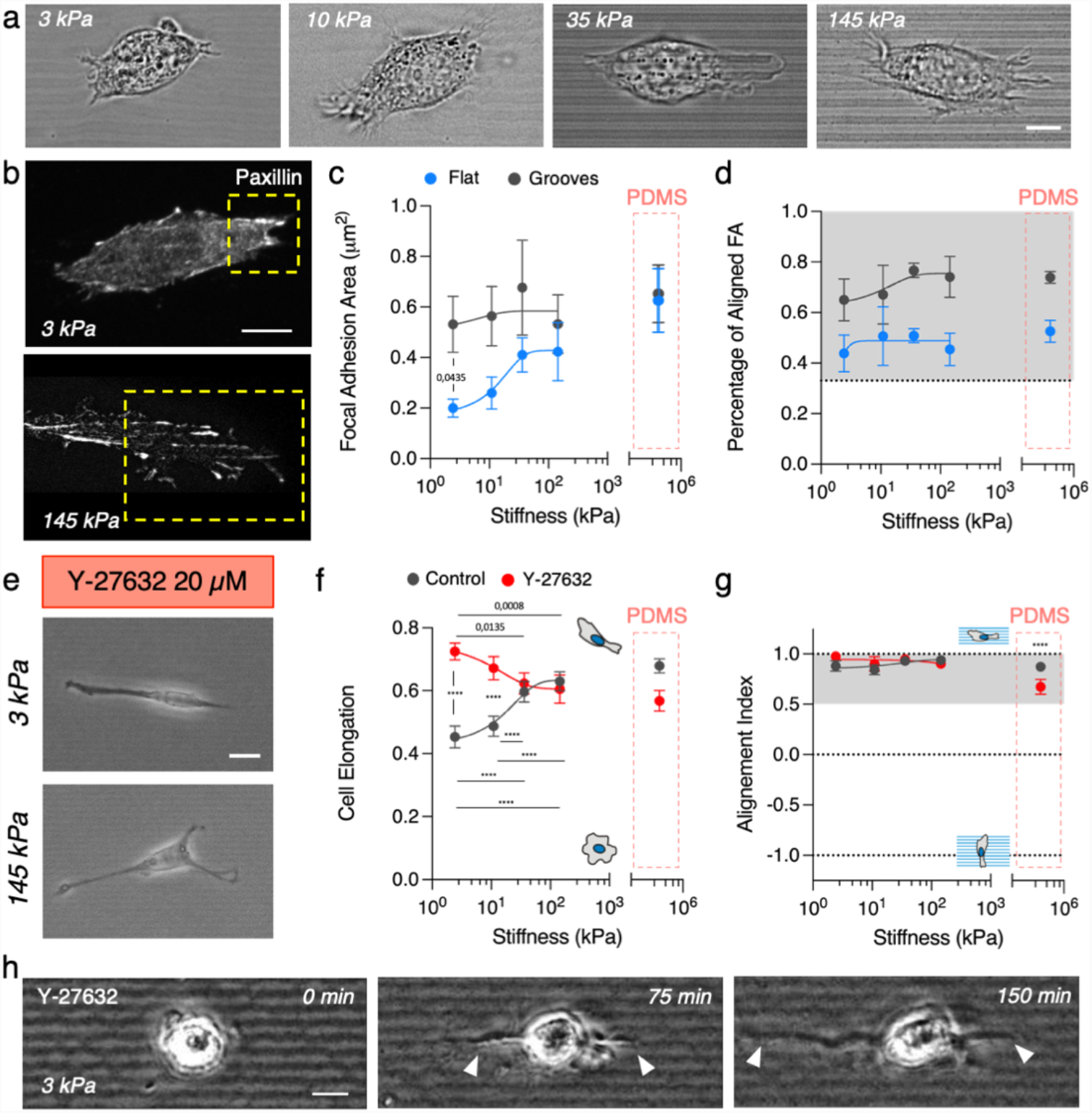
Decrease of cell contractility increases contact guidance on soft topographical grooves and ridges. (**A**) DIC images of fibroblasts on microsctructured PAA gels. Scale bar, 8 μm. (**B**) Paxillin immunostaining of fibroblast on grooves of 3 kPa and 145 kPa. Examples of lamellipodial regions used for focal adhesion analysis is highlighted in yellow. Scale bar, 10 μm. (**C**) Focal adhesion area and (**D**) percentage of aligned focal adhesions as a function of increasing substrate stiffness. Grey area in (**D**) corresponds to values above the expected for a random distribution. (**E**) Representative phase contrast images of 3T3 fibroblast treated with 20 μM Y-27632 on 2 μm wide grooves. Scale bar, 20 μm. (**F**) Cell elongation and (**G**) cell alignment index as a function of the substrate stiffness. (**H**) Phase contrast images of a NIH 3T3 fibroblast treated with 20 μM Y-27632 spreading on 2 μm wide grooves of 3 kPa PAA. Scale bar, 10 μm. See Tables S3 and S4 for the number of cells and experiments. Data points (Mean ± SE) in (**C**) and (**D**) and (Mean ± CI) in (**F**) and (**H**) were fitted as an eye-guide. Statistical significance was assessed by Tukey’s tests (**F** and **G**), Welch’s test (**C**) and Kruskall-Wallis’ test (**D**).

Onto stiff substrates, it is established that focal adhesions align and mature along the groove direction, leading to the alignment of associated stress fibers, and generating anisotropic forces that translate into cell alignment (*33*), therefore linking topographical contact guidance to cell contractility (*21, 28*). In this scenario, decreasing NIH 3T3 fibroblasts contractility through the inhibition of the Rho-associated protein kinase (ROCK) led to a decrease of cell elongation and alignment when cultured on stiff PDMS grooves (Fig. 2f,g). We then performed an analogous set of experiments but culturing NIH 3T3 fibroblasts onto soft PAA substrates. Strikingly, a different response was observed on these materials. On the stiffest gels (35 – 145 kPa) cell elongation remained unchanged, but cells on the softest gels (3 – 10 kPa) showed and increased elongation despite fibroblast contractility was decreased (Fig. 2e,f). However, this change in morphology had little impact on cell alignment, which, contrary to what was observed in PDMS substrates, was preserved (Fig. 2g). Although cell contractility is necessary for contact guidance on stiff substrates, initial cell polarization through contact guidance has been found to be contractility independent (*22*). Then, we argued that our results on soft gels may be biased by the cells being already polarized when the ROCK inhibitor was added. To clarify this point, we further repeated the experiments seeding the cells onto 3 kPa grooves in the presence of the ROCK inhibitor. We then observed that initially roundish fibroblasts formed long protrusions emerging from opposite sides of the cell body and that these protrusions extended following the direction set by the grooves (Fig. 2h), leading to larger cell elongation than non-treated cells. This result confirms that fibroblast contractility is involved in regulating contact guidance onto soft gels (3 – 10 kPa) and that decreasing cell contractility allows cells to further elongate, thus increasing their contact guidance for these soft substrates. Taken all these observations together, the biphasic behavior found in the response of topographical contact guidance to changes in cell contractility for soft and stiff substrates seems to indicate that different molecular actors for mediating the response to topographical patterns may be involved as a function of the substrate rigidity.

### Microtubule polymerization mediates topographical contact guidance of fibroblast in a stiffness dependent manner

We found the phenotype of ROCK-inhibited fibroblasts reported above as being reminiscent of the growth cones of neurites, which are cellular structures rich in actin and microtubules (*23, 48*). We then hypothesized that actin, but also microtubules may be involved in mediating the cellular response to topographical patterns for different substrate rigidities. To investigate this point, we explored the actin and tubulin distribution on fibroblast cultured on grooves of different stiffness. Our results show that F-actin structures filled in the grooves for all of substrates tested (Fig. 3a). On the contrary, microtubules, stained with *α*-tubulin, showed a progressive increasing conformity with the grooves and ridges when increasing the PAA substrate stiffness (Fig. 3a). Noteworthy, the geometry of the grooves, which was revealed in the fluorescence signal captured at the basal plane of the cells, lost definition when decreasing substrate stiffness. This effect could be linked to the pattern deformation due to cell contractility (Fig 2a). Despite this deformation, when averaging actin intensity profiles across several cell lamellipodia *vis-à-vis* the grooves, we observed actin conformity with them for all the PAA gels (Fig. 3b,c). Strikingly, tubulin intensity profiles showed that in-groove microtubule signal exhibited no conformity with topography on 3 kPa substrates, while it was evident on > 35 kPa substrates (Fig. 3d). This behavior of the microtubular network can be related to a stiffness-dependent microtubule acetylation which stabilizes the microtubular network (*49, 50*). Thus, these results point out the potential role of microtubules on the cell response to topographical patterns in a stiffness dependent manner.

**Fig. 3.**
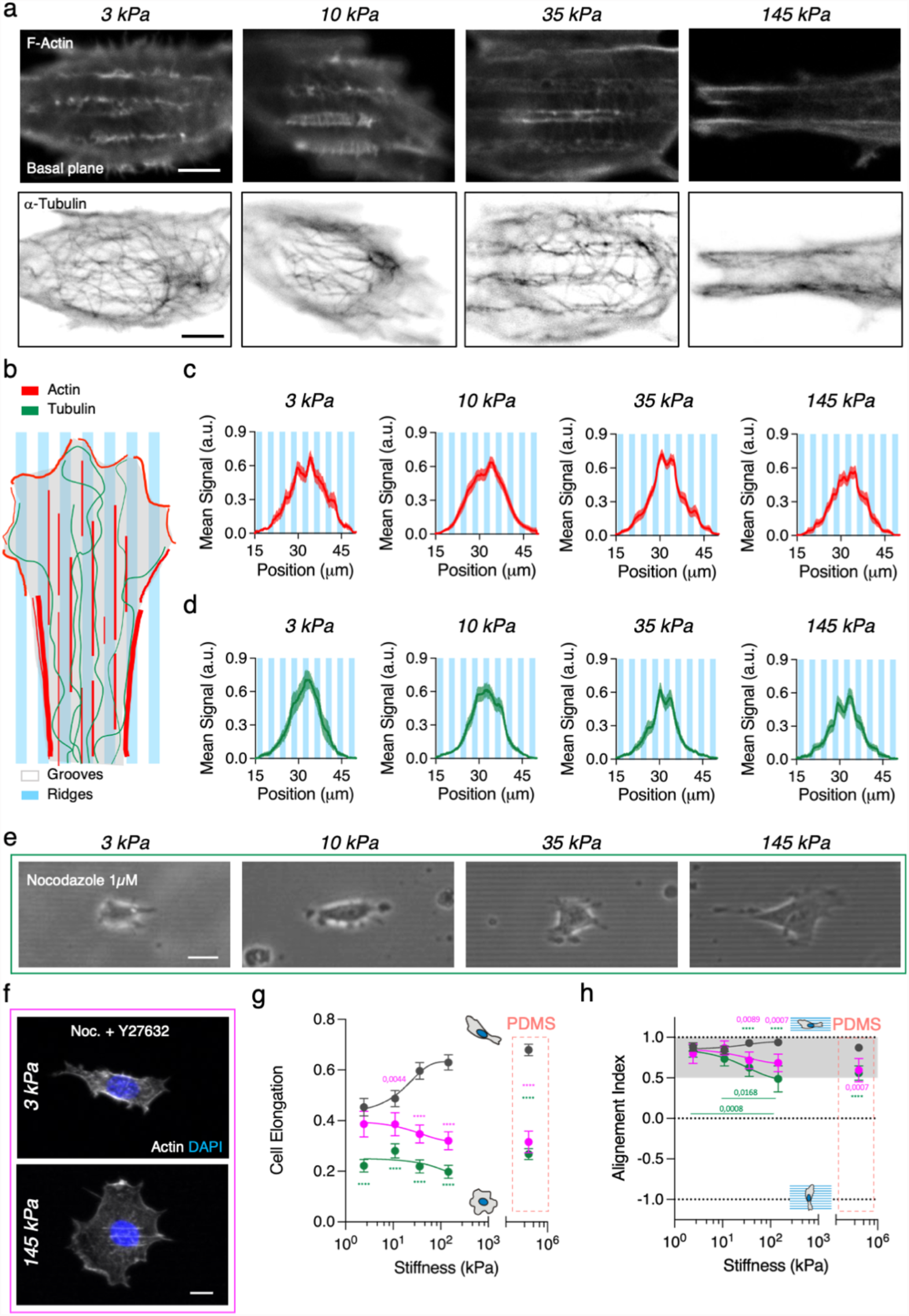
Microtubule polymerization mediates NHI 3T3 fibroblasts response to topographical patterns. (**A**) F-actin (phalloidin) immunostaining of fibroblasts on grooves of increasing stiffness. Scale bar, 5 μm. *α*-tubulin immunostaining of fibroblasts on grooves of increasing stiffness. Scale bar, 5 μm. (**B – C**) Average intensity profile of LifeAct-GFP signal across the grooves and ridges at the lamellipodial region for increasing substrate stiffness. Mean ± S.E.M. (**D**) Average intensity profile of SiR-tubulin signal across the grooves and ridges at the lamellipodial region for increasing substrate stiffness. Mean ± S.E.M. (**E**) Representative phase contrast images of 3T3 fibroblast treated with 1 μM nocodazole on 2 μm wide grooves of increasing stiffness. Scale bar, 20 μm. (**F**) Representative images of 3T3 fibroblast treated with 20 μM Y-27632 and 1 μM nocodazole on 2 μm wide grooves. Scale bar, 10 μm. (**G**) Cell elongation and (**H**) cell alignment index as a function of the substrate. See Tables S5 – 8 for the number of cells and experiments. Data points (Mean ± CI) in (**G**) and (**H**) were fitted as an eye-guide. Statistical significance was assessed by Tukey’s tests (**H** and **G**).

Next, to further investigate the involvement of microtubules when considering substrate rigidity on contact guidance-associated phenomena, we used Nocodazole to depolymerize microtubules. Then, we evaluated the effects of this drug on topographical contact guidance hallmarks evidenced by cell morphology (cell area, elongation and alignment) on grooved substrates. Nocodazole treatment led to uncoordinated protrusion dynamics (Fig. 3e), and while cell area was roughly maintained, cells became less elongated and lost the elongation stiffness-dependency (Fig. 3g). Concomitantly with the decrease in cell elongation, nocodazole treatment produced a progressive misalignment between the cells and the grooves as substrate stiffness increased (Fig. 3h). This progressive misalignment followed the inverse trend of the progressive stiffness-dependent tubulin conformity with the grooves and ridges observed for non-treated cells. Taken together, these results seem to point out that microtubules are indeed involved in a stiffness-dependent cell response to topographical contact guidance.

Microtubule depolymerization enhances cell contractility through an increase of RhoA activity (*51*). Thus, results described above are a convolution of microtubule depolymerization and contractility enhancement. To disentangle both effects, we used ROCK inhibition to decrease cell contractility in nocodazole treated cells (Fig. 3f). This reverted the increase in contractility induced by microtubule depolymerization. As a result, cells were more elongated than when treated with Nocodazole alone (Fig. 3g). On the softest grooves (3 kPa), cells retained an elongation similar to non-treated (control) cells, but they progressively became less elongated than their non-treated counterpart when increasing substrate stiffness. In addition, and despite the decrease in cell elongation with the stiffness of the substrate, cells showed actin structures partially aligned with the grooves for stiffness > 10kPa (Fig. 3f and Fig. S3a). This shows that cells can elongate through contact guidance in the absence of microtubules on soft substrates, but that the stiffer the substrate becomes, the more the cells rely on the microtubule cytoskeleton to regulate elongation, and that it cannot be counterbalanced by the topographical pattern. These actin structures co-localized with accumulations of the focal adhesion protein paxillin (Fig. S3b). The quantitative evaluation of the focal adhesions showed that they were smaller and less aligned with the grooves than in non-treated (control) cells (Fig. S3c-d). This, in turn, translated into a decrease of cell alignment with the grooves when increasing stiffness (Fig. 3h). While cells on 3 kPa and 10 kPa substrates aligned with the grooves an amount similar to non-treated cells; on stiffer substrates cell alignment was significantly diminished. Thus, taken together, these results show that microtubule polymerization mediates cell elongation and alignment by topographical contact guidance in a stiffness-dependent fashion.

### Microtubules are stiffness-dependent key regulators for fibroblast effective migration by topographical contact guidance

Cell elongation and alignment such as those observed here lead to directed cell migration on stiff substrates. Therefore, we next checked if this phenomenon was also observed on our grooved substrates of different rigidities. For this purpose, cell motility was tracked by time-lapse microscopy. The results obtained showed cell’s trajectories taking place in a narrow spatial distribution confined along the direction axis set by the grooves for all the stiffness studied (Fig. 4a and Movie S2). We then measured the total distance covered by the cells, their net displacement, and the degree of directionality of their instantaneous movements (Fig. 4b, see Methods section for details) during 6 hours of migration. The total cell migration distance increased with substrate stiffness until reaching a plateau above ∼35 kPa for both flat and grooved substrates (Fig. S4). On the contrary, the net cell displacement was enhanced by the presence of the grooves and was also increased with the substrate stiffness (Fig. 4c). Additionally, in contrast to flat substrates (directionality index ∼ 0), grooves set a directional cell migration (directionality index > 0.5) for all the substrates tested (Fig. 4d). Altogether, our results show that for fibroblasts the directed cell migration associated to topographical contact guidance is effective across several orders of magnitude in substrate stiffness. Importantly, this range comprises, aside from the well-studied non-physiological range, physiologically and pathologically relevant stiffness values.

**Fig. 4.**
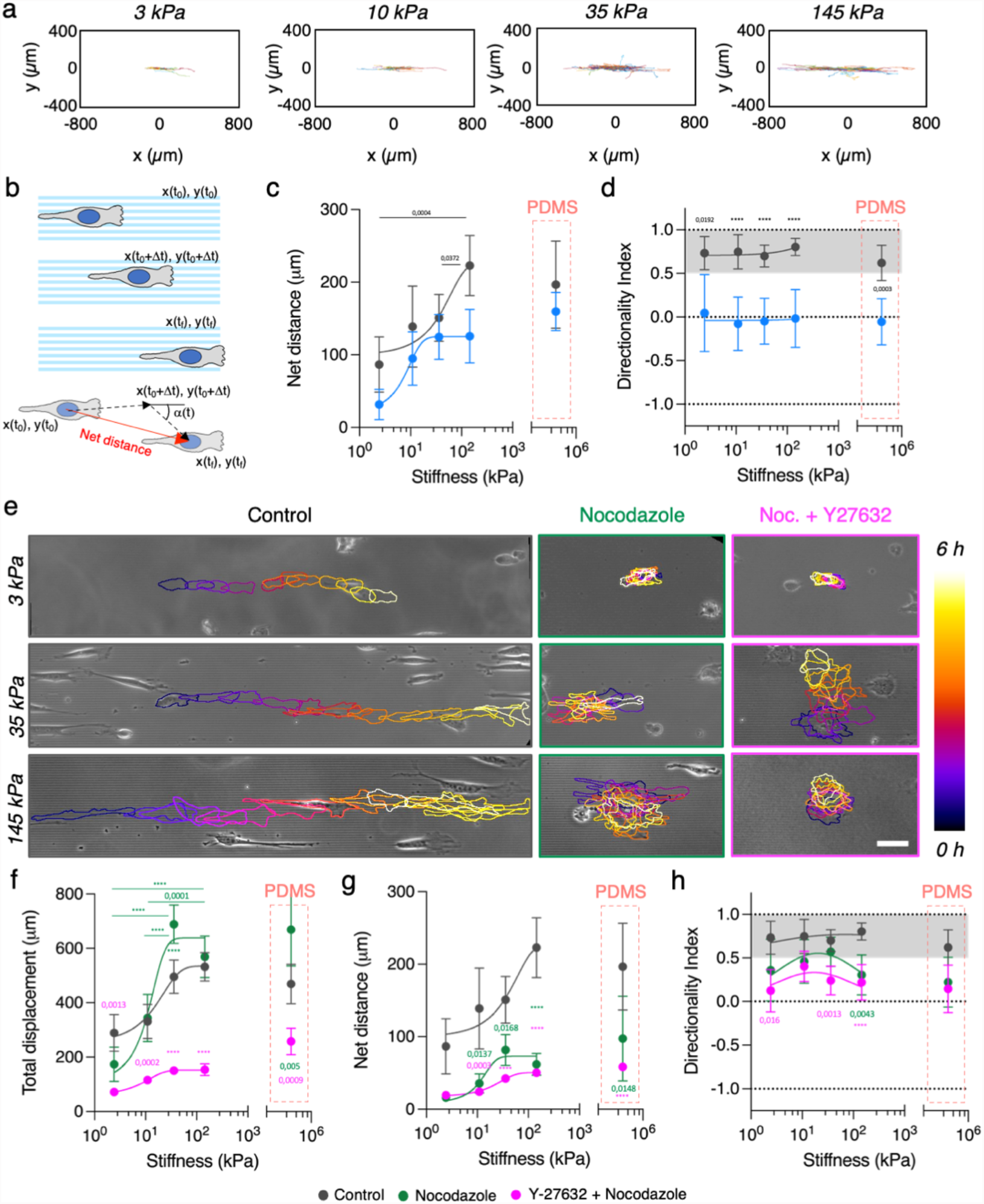
Microtubule depolymerization leads to a stiffness dependent topographically guided migration. (**A**) Single cell trajectories on grooved hydrogels of different stiffness. (**B**) Scheme depicting the quantification of cell trajectories. (**C**) Cells’ net distances versus substrate stiffness. (**D**) Trajectory directionality as a function of substrate stiffness. (**E**) Cell trajectories on grooves of different stiffness for control cells, cells treated with nocodazole and cells treated with Y27632 and nocodazole. Cell outlines are color-coded by time. Scale bar, 50 μm. (**F**) Total displacement, (**G**) net distance and (**H**) trajectory directionality index of cells. See Tables S10 – S12 for the number of cells and experiments. Data points (Mean ± CI) in (**C**), (**D**), and (**F** – **H**) were fitted as an eye-guide. Statistical significance was assessed by Tukey’s tests.

So far, we observed that directed migration by topographical contact guidance happened independently of substrate stiffness, but we did indeed observed differences in the role of microtubules in contact guidance in a stiffness-dependent manner. Thus, we decided to explore further the cytoskeletal actors involved in directed migration through contact guidance and check whether the stiffness dependent effects of microtubule depolymerization on cell shape and alignment were also present in the directed cell migration. To do that, we tracked the movements of cells with microtubules depolymerized and increased contractility (nocodazole treatment) and with depolymerized microtubules (nocodazole and Y27632 ROCK inhibitor treatment). Results showed that cell migration was in fact severely affected by the lack of microtubule polymerization (Fig. 4e and Fig. S5). Despite nocodazole treatment did not greatly change cell total displacement (Fig. 4f), the cell net migration distance decreased drastically for all the stiffnesses studied (Fig. 4g). Upon nocodazole treatment cells experience a directional switch effect (*52*) that, along with uncoordinated protrusions, can lead to non-directed migration and poor net motion. Indeed, nocodazole-treated cells migrated non-directionally on 145 kPa PAA grooves (Fig. 4h). On the contrary, a less persistent but directed migration was observed on softer grooves (10 – 35 kPa) (Fig. 4h), evidencing that the role of microtubules is less relevant in leading to directed migration by contact guidance for soft substrates and, thus, suggesting that substrate stiffness modulates the contribution of the cytoskeletal players to the contact guidance outcome. On the other hand, when the increase in contractility caused by nocodazole treatment was reverted with ROCK inhibitor, cell migration was significantly reduced both in terms of total displacement and net distance for all the substrates tested (Fig. 4f,g and Fig. S5). Moreover, directionality was now further decreased compared to nocodazole-treated cells (Fig. 4h). In this case, the inhibition of both cell contractility and tubulin polymerization led to the loss of directional migration when compared to non-treated cells for soft (3 kPa), intermediate (35 kPa) and stiff (145 kPa) substrates (Fig. 4h). Altogether, these results suggest that, despite directed migration induced by contact guidance takes place for all the range of substrate stiffness in fibroblasts, it involves multiple cytoskeletal actors whose relative contributions may change with stiffness, with acto-myosin contractility being more relevant on soft substrates and microtubules becoming more relevant as substrate rigidity increases.

### T47D carcinoma cells perform directed migration induced by contact guidance in a stiffness dependent manner

Up to this point, our experiments revealed that mesenchymal-like cells experienced topographical contact guidance and directed cell migration regardless the stiffness of the substrate, even if complex stiffness-dependent mechanisms are involved in the process. Moreover, we also saw that epithelial cells also experienced topographically induced elongation and alignment, but this was strongly dependent on the substrate stiffness. Therefore, we wondered if for epithelial cells the substrate stiffness affected directed cell migration. To check this, we seeded T47D carcinoma cells at low density and the motility of individual cells was tracked by time-lapse microscopy. In agreement with previous experiments (*33*), on stiff PDMS substrates T47D cells also experienced directed migration by contact guidance but their trajectories were much shorter than the ones observed for 3T3 fibroblasts (Fig. S6). Noteworthy, when T47D epithelial cells were cultured on softer substrates (approaching physiological and pathological values of soft tissues), single cell trajectories were significantly affected by the grooves rigidity and differed greatly from substrates with non-physiological stiffness values. Specifically, cell trajectories were longer and became more restricted to the direction set by the grooves than on stiff PDMS (Fig. 5a, Movie S3). In addition, total cell displacement, net distance, and directionality index increased when topographical patterns became stiffer (up to 145 kPa). Basically, these parameters showed that directed migration by contact guidance for T47D cells had a non-monotonic behavior with the substrate stiffness within the range tested (Fig. 5b – d), being increasingly effective for cells cultured on physiological stiffness but decreasing for cells cultured on non-physiological ones. Epithelial cells form focal adhesions that tend to be small, their microtubules are organized in setting cell apical-basal polarity, and they do not migrate directionally as single cells in physiological conditions (*45*). Remarkably, in here we have found that non-migratory T47D cells can develop a stiffness-dependent migratory phenotype when cultured on topographical grooves with rigidities similar to the ones encountered on soft tissues, highlighting the relevance of including the physiological values of the matrix when addressing topographical contact guidance for this cell type.

**Fig. 5.**
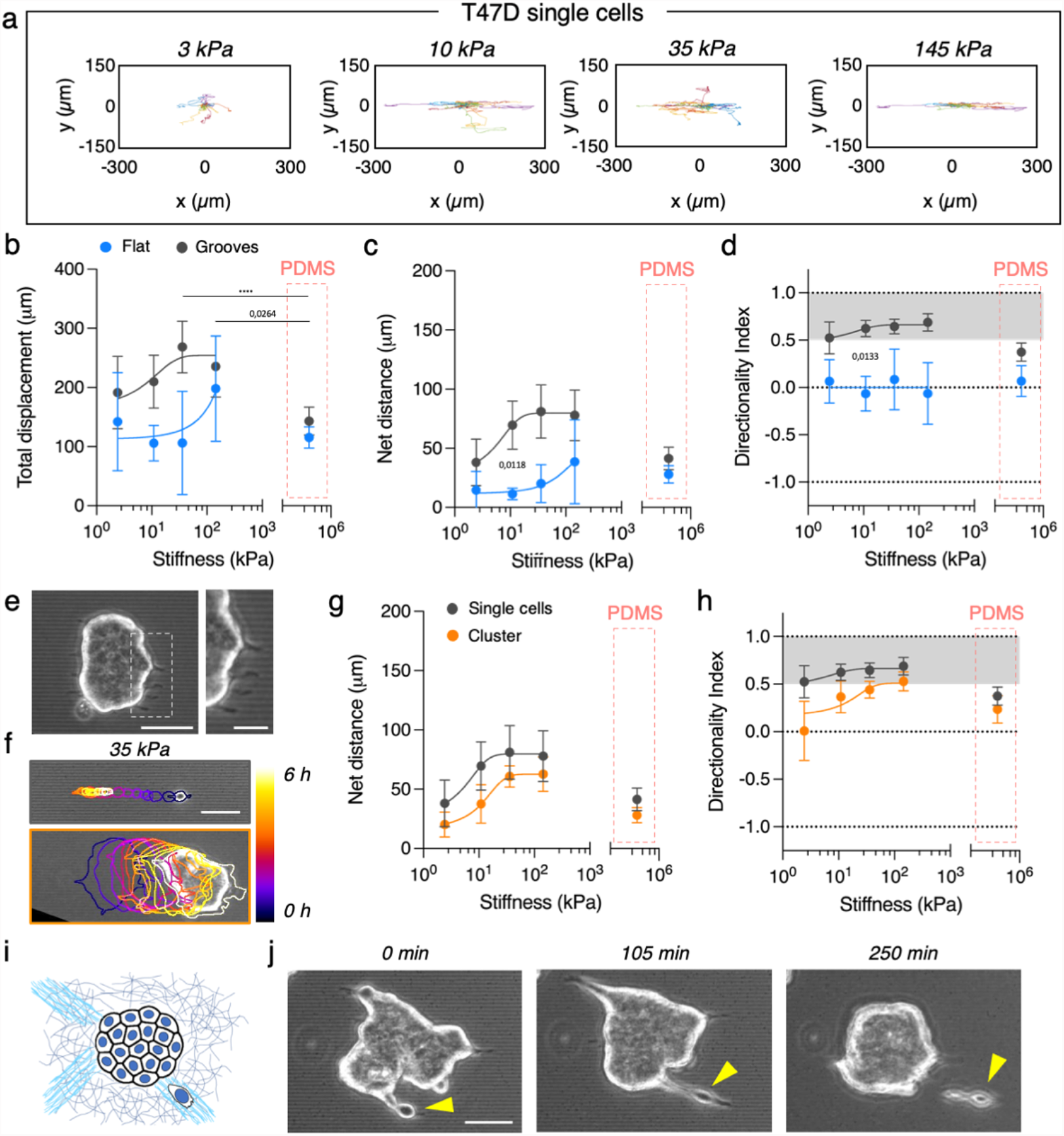
T47D carcinoma cells perform directed migration as a function of stiffness. (**A**) Single T47D cells’ trajectories on grooves of increasing stiffness. (**B**) Total displacement, (**C**) net distance and (**D**) trajectory directionality index of single T47D cells migrating on flat and grooved polyacrylamide hydrogels of increasing stiffness. (**E**) Clusters of T47D cells extend protrusions along the grooves. Scale bar, 50 μm. Inset scale bar, 20 μm. (**F**) Cell trajectories on grooves of 35 kPa for control single T47D cells and T47D cell clusters. Outlines are color-coded by time. Scale bar, 50 μm. (**G**) Net distance and (**H**) directionality as a function of groove stiffness for T47D cell clusters and single T47D cells. (**I**) Scheme of a cluster of T47D cells and single T47D cells navigating on aligned extracellular matrix fibers. (**J**) Snapshots of a T47D cell detaching from the periphery of a cell cluster. See Tables S13 for the number of cells and experiments. Data points (Mean ± CI) in (**B – D**), and (**G** – **H**) were fitted as an eye-guide. Statistical significance was assessed by Tukey’s tests.

Finally, T47D epithelial cells formed medium-sized clusters as well (Fig. 5e) that allowed us to investigate directed collective migration by contact guidance and its dependence on the stiffness of the substrate. When in the presence of topographical patterns, we observed that T47D clusters formed cell protrusions along the grooves direction (Fig. 5e inset) and in some cases performed directed migration (Fig. 5f, Fig. S7 and Movie S4). Similar to single cells, T47D clusters barely migrated on PDMS topographical patterns, and their movement was close to random (Fig. 5g,h). However, when seeded on softer topographical patterns, they did show a larger motility. Again, a non-monotonous trend with the substrate stiffness was observed for the effective migration of the clusters and their net distance (Fig. 5g,h). On soft 3 kPa grooves, clusters migrated randomly. Then, this migration became progressively more directed when the stiffness of the underlying substrate increased (Fig. 5h), before decreasing in directionality again for very hard substrates (PDMS). Taking these data together, we observed that epithelial cells both as single cells and as cell clusters perform directional and efficient migration triggered by topographical contact guidance when they are presented with substrates that possess stiffness values within the physiological range of soft tissues.

## Discussion

Contact guidance has been known for more than a century (*53*), but the key aspects of how cells sense the underlaying topographical patterns and how the cytoskeletal elements contribute to this phenomenon have started to be clarified only in recent years and are still being scrutinized (*33, 34, 54*). Mechanics plays a central role in the behavior of the cellular structures involved in sensing contact guidance, but the mechanical properties of the substrates have been completely overlooked when studying this phenomenon. Only recent developments in microfabrication techniques have allowed to pattern soft gels (*42, 55*) and study contact guidance in the range of stiffness relevant for soft tissues. For example, T-cells perform topographically induced contact guidance only onto soft grooves (16 kPa) (*40*). In here we show that actually the stiffness of the topographical patterns modulates the contact guidance response of fibroblasts and epithelial cells (Fig. 1). Grooves enhance the elongation of ‘front-back’ polarized fibroblasts across all the stiffness range, being the highest impact observed on soft substrates (3 kPa). On the contrary, grooves can break symmetry and elongate epithelial cells, which typically have apical-basal polarization, when the substrate stiffens (35 – 145 kPa).

These changes in contact guidance as a function of the rigidity of the topographical grooves involve several cytoskeletal actors. While focal adhesion alignment and actin conformity with the topographical shapes were observed for all the stiffness range (Fig. 2d and Fig. 3a,c), the response of the microtubule cytoskeleton to the topographical pattern was stiffness-dependent (Fig. 3a,d). Further inhibition of tubulin polymerization showed that the loss of both cell alignment and directional migration increased with substrate stiffening (Fig. 3h and Fig. 4h). On the contrary, cell elongation and alignment to the grooves were invariant to tubulin depolymerization on soft substrates but were sensitive to changes in contractility (Fig. 2f and Fig. 3g). These results suggest that there is a balance between cell contractility and tubulin polymerization in regulating fibroblasts response to topographical grooves. Furthermore, when both actomyosin contractility and tubulin polymerization were inhibited, this hindered the ability of the focal adhesions to align with the grooves and ridges. Whether this effect was due to a lack of cell contractility (*33*) or to the role of microtubules in the dynamics of focal adhesion complexes during mechanosensation (*50, 56, 57*) remains an open question.

In a previous study, perturbation of the microtubule-contractility axis regulated T-cell contact guidance when groove elasticity changed (*40*). Here, we show that in the case of fibroblasts actomyosin contractility regulates contact guidance on soft substrates as well, but microtubules take over in being the major regulator of contact guidance when the substrate stiffens. This translated into a stiffness independent contact guidance migration for fibroblasts, which was reported for T-cells only when Rho-A activity was increased through tubulin depolymerization. This highlights the importance of the cell phenotype (and the associated cytoskeleton organization) on experiencing contact guidance. We could show that in breast carcinoma epithelial cells, substrate stiffness modulated the cell’s response to the topographical patterns in a third manner: contact guidance increases as the groove rigidity increases (Fig. 1 and Fig. 5). Unlike fibroblasts, epithelial cells have small focal adhesions and apico-basal polarized microtubule networks (*45*), so the interaction between adhesions, actomyosin and microtubules unfolds with stiffness differently than for fibroblasts. This effect was even more dramatic when cell-cell adhesions were involved. Epithelial clusters barely moved on PDMS, but when substrate stiffness was decreased to more physiological values, they migrated collectively in the direction of the grooves. This behavior was again abrogated when the stiffness was further decreased. In our experiments, topographical patterns with intermediate stiffness (∼35 kPa) could be viewed as stiff and aligned protein fibers of the tumoral stroma (Fig. 5i). Cells at the periphery of tumor cell clusters may use these stiff aligned fibers to adhere, align and migrate outwards, detaching from the cluster (Fig. 5j). This shows that both matrix stiffening (*58*) and alignment of extracellular matrix fibers (*10*) could synergize during metastasis and cancer cell invasion. Moreover, since stiffness-dependent contact guidance was also observed for cell clusters, not only single cells but cell clusters could take advantage of aligned and stiff fibers during the process of invasion.

Together, our findings demonstrated that substrate stiffness mediates cell response to topographical grooves. We showed that mesenchymal-like cells relay on different cytoskeletal actors to mediate their response to their environment’s architecture depending on substrate stiffness. Actomyosin contractility on soft grooves and tubulin polymerization on stiff ones cooperate to produce a stiffness independent output to contact guidance. Moreover, we demonstrated that an increase in groove stiffness enhanced directional response of epithelial cells in a biphasic fashion. And that epithelial-like cells experience stiffness dependent contact guidance even as multicellular clusters, where cell-cell adhesions are involved. Therefore, only acknowledging substrate stiffness when studying the effects of topography in cell behavior, we will be able to fully comprehend how cells respond to anisotropic matrix architectures *in vivo*.

## Materials and Methods

### Experimental design

To explore the impact of substrate stiffness on the ability of topography to influence cellular alignment and contact guided migration, we cultured mesenchymal and epithelial cells on 2 μm wide and 1 μm deep topographical grooves and ridges made of PAA gels of different stiffness (3 – 145 kPa). Detailed steps are described below.

### Cell culture

NIH 3T3 mouse embryonic fibroblasts and T47D cells were obtained from ATTC. NIH 3T3 fibroblasts were grown with DMEM (Invitrogen) medium supplemented with 10% Fetal Bovine Serum (FBS) (Life Technologies), 1% L-glutamine (Gibco), 1% penicillin-streptomycin (Sigma-Aldrich), 1% sodium pyruvate (Invitrogen) at 37ºC, and 5% CO2. Cells were passaged every 2 – 3 days. T47D cells were grown with RPMI-1640 medium (Invitrogen) supplemented with 10% FBS at 37ºC, and 5% CO2. Cells were passaged every 2 – 3 days. For the experiments employing NIH 3T3, 50000 cells per well (35 mm petri dish) were cultured on the polyacrylamide (PAA) gels during 24 h. For the experiments employing T47D, 75000 cells per well (35 mm petri dish) were cultured on the PAA gels during 24 h.

### Polyacrylamide gel fabrication and microstructuring

PAA microstructured gels were prepared following the experimental procedure described in (*42*). Briefly, we first prepared PDMS (Sylgard 184 Silicon Elastomer, Dow Corning) molds (5 mm in diameter containing 2μm width and 1 μm high grooves) and spacers (10 mm in diameter). Then, 35 mm bottom glass petri dishes (MatTek) were silanized with 3-(aminopropyl)trimethoxysilane (Sigma-Aldrich) following the procedure described in (*59*). Afterwards, the PDMS spacer was placed on the silanized glass of the petri dish, filled with 70 μL of polyacrylamide solution (see Supplementary Table S14) and covered with a poly(ethylene naphtalate) (PEN) (Goodfellow) sheet with the PDMS mold attached. PAA gels were then kept at room temperature (RT) with a small weight on top of the PEN sheet for 2 h to polymerize. The polymerized gels were demolded by carefully removing the flexible mold and then stored in PBS at 4ºC until further use, allowing them to achieve equilibrium swelling.

Before cell seeding, fibronectin from bovine serum (Sigma-Aldrich) was covalently bound to PAA gels using Sulfo-SANPAH reagent (Sigma-Aldrich). PBS was removed and then, we freshly prepared a Sulfo-SANPAH solution at a concentration of 2 μg/μL in Milli-Q water and added 50 μL on top of each hydrogel. The PAA gels with the Sulfo-SANPAH solution were UV-irradiated (Light Source LQ-HXP 120 UV, LEJ) for 60 s and then rinsed several times with PBS. Next, we incubated the proteins on the activated gels using 50 μL of a 20 μg/mL fibronectin solution for 1 h at RT. Functionalized gels were stored in culture medium until cell seeding.

### Immunostaining

Cells were fixed with 10% neutralized formalin (Sigma -Aldrich) for 35 min at 4ºC and washed three times with PBS. Cells were then permeabilized with 0.5% Triton X-100 (Sigma) for 30 min and blocked for at least 2 h with a blocking buffer containing 1% BSA (Sigma), 3% donkey serum (Millipore), and 0.2% Triton X-100 in PBS. All samples were incubated with the primary antibodies overnight at 4°C, followed by a 2 h incubation at RT with secondary antibodies, and phalloidin rhodamine (, Cytoskeleton) to stain filamentous actin (F-actin) (1:140). Finally, 30 min incubation with 4′,6-diamidino-2-phenylindole (DAPI, Life Technologies) (1:1000) for nuclei staining was employed. After three PBS washes, samples were kept in Fluoromount-G mounting media (SouthernBiotech) and stored in a humid chamber at 4ºC until imaged.

### Antibodies

Imaging of actin cytoskeleton and microtubule network in life cell experiments was performed using NIH 3T3 stable cell line expressing Lifeact-GFP (Addgene) treated with SiR-tubulin (Spirochrome). NIH 3T3 fibroblasts were transfected with Lifeact-GFP using Lipofectamine 2000 (Life Technologies) and selected using geneticin G418 antibiotic (ThermoFischer). Fibroblasts expressing Lifeact-GFP were seeded on the PAA gels and cultured for at least 12 h. Then, cell culture media was replaced by fresh media containing 100 nM SiR-tubulin and incubated for 12 h. After that, cells were imaged under the microscope at 37ºC in a humidified atmosphere containing 5% CO2.

### Image acquisition

Phase contrast images were acquired using a Nikon Eclipse Ts2 with x10 (ADL Ph1 0.25 NA) and x20 (ADL Ph1 0.40 NA) objectives. Migration experiments were performed by using phase contrast microscopy in a Axio Observer 7 (Carl Zeiss) using a x10 objective for 6 h at 37ºC and 5% CO2. Life cell imaging of actin and microtubules was performed in a Axio Observer 7 (Carl Zeiss) using a x63 objective. Immunostained samples were observed using confocal laser scanning microscope (LSM 800, Zeiss) equipped with a x63 oil objective (NA 1.40) and a x100 oil objective (NA 1.40).

### Cell morphology evaluation

The effects of both stiffness and topography were established by measuring morphological parameters using Image J software (http://rsb.info.nih.gov/ij, NIH). Cell area was measured by outlining the cell shape. Cell elongation and alignment were measured by fitting an ellipse to the cell shape. The elongation was assessed by calculating (*a* − *b*) / (*a* + *b*), being a and b the ellipse major and minor axis, respectively. The alignment index was calculated as *cos* (2 · *θ*) where θ is the angle between the ellipse’s major axis and the direction of the topographical pattern (Figure 1c).

### Cell migration evaluation

The centroid trajectories of NIH 3T3 cells and T47D single cells were tracked using the Manual Tracking Plug-in in Fiji (http://rsb.info.nih.gov/ij, NIH, USA). For T47D clusters, each cluster was outlined and its centroid was calculated with Fiji. Data analysis was performed using a custom-made code in Matlab (Mathworks, USA). Cell centroid positions during the experiment were defined as (*x*(*t*_*i*_), *y*(*t*_*i*_)), being *t*_*i*_ = *i* · *Δt* with *i* = 0, …, *n* and *Δt* the time between consecutive images. The total distance covered by the cells was computed as 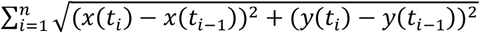 and the net distance as the vector difference between the initial and the final point 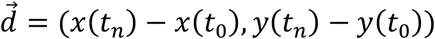, being its module 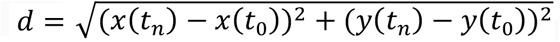. The alignment of each trajectory was computed by averaging the alignment of the vector displacement between two consecutive time-points and the topographical pattern for every time-point ⟨cos (2 · *α*(*t*_*i*_)⟩_*i*_, being *α*(*t*_*i*_) the angle between the vector defined by two consecutive cell positions (*x*(*t*_*i*_) − *x*(*t*_*i*−1_), *y*(*t*_*i*_) − *y*(*t*_*i*−1_)) and the direction of the grooves. This index equals 1 when the trajectory is parallel to the direction of the grooves, 0 when is random and -1 when it is perpendicular.

### Focal adhesion analysis

To characterize focal adhesion size, shape and orientation, fluorescent images of paxillin were treated using Fiji as follows. First, a median filter was applied and the background signal subtracted. Then, a contrast-limited adaptive histogram equalization (CLAHE) was used and the background was again removed using an exponential function. Finally, a Laplacian of Gaussian filter was applied to the resulting image and the focal adhesions were selected with auto-threshold function. Orientation of the focal adhesions was measured *vis-à-vis* the long cell axis for flat gels and *vis-à-vis* the grooves direction for topographical patterns.

### Actin and tubulin intensity profiles

To obtain actin and tubulin conformity with the topographical patterns, fluorescent images of live cells expressing Lifeact-GFP and treated with SiR-tubulin were used. Intensity profiles were acquired across the cell lamellipodia using the line function in Fiji. A line 11 μm wide perpendicular to the topographical pattern was used. Single intensity profiles were individually normalized by their maximum intensity. Then, the average for several cells was computed.

### Chemical treatments

To inhibit ROCK activity, 20 μM Y-27632 (Sigma-Aldrich) was added to the cell culture media. To disrupt microtubules and inhibit microtubule dynamics we added 1 μM nocodazole (Sigma-Aldrich) to the cell culture media. Drug treatments were added at least 6 h after cell seeding, were incubated for at least 6 h before measurements were performed and were maintained until the end of the experiment.

### Statistics

No statistical methods were used to predetermine sample size. Measurements were performed on individual cells or cell clusters (n) obtained in different independent experiments (N). The exact values are defined in Supplementary Tables I – XIV. Data presentation (as Mean value ± standard deviation (SD), Mean value ± 95% confidence interval (CI) or as Mean value ± standard error of the mean (SE)) is defined at the corresponding figure caption. Data fitting was performed using GraphPad Prism 9 to serve as an eye-guide. The statistical test were performed using GraphPad Prism 9 and are defined at the corresponding figure caption. The p-values are specified in the figures and when they are not mentioned, differences are not statistically significant (p > 0.05).

## Acknowledgments

We thank the Martinez Lab members and P. Roca-Cusachs for discussions and help, and A.L. Godeau and R. Sunyer for critical reading of the manuscript.

Funding for this project was provided by:

European Union Horizon 2020 ERC grant (agreement no. 647863 -COMIET)

CERCA Programme/Generalitat de Catalunya (2017-SGR-1079)

Spanish Ministry of Economy and Competitiveness (TEC2017-83716-C2-1-R, and the Severo Ochoa Programme for Centres of Excellence in R&D 2016-2019)

The results presented here only reflect the views of the authors; the European Commission is not responsible for any use that may be made of the information it contains.

## Author contributions

Conceptualization: JC, EM

Methodology: JC, EM

Investigation: JC, VF-M, VA, BR-C

Validation: JC

Formal Analysis: JC

Visualization: JC

Supervision: JC, EM

Writing—original draft: JC, EM

Writing—review & editing: JC, VF-M, VA, BR-C, EM

Project administration: JC

Funding acquisition: EM

## Data and materials availability

All data are available in the main text or the supplementary materials.

