## Supplementary Material for "Soft topographical patterns trigger a stiffness-dependent cellular response to contact guidance"

**This PDF file includes:**

Figs. S1 to S7
Tables S1 to S14
Movies S1 to S4

**Other Supplementary Materials for this manuscript include the following:**

Movies S1 to S4

**Fig. S1.**

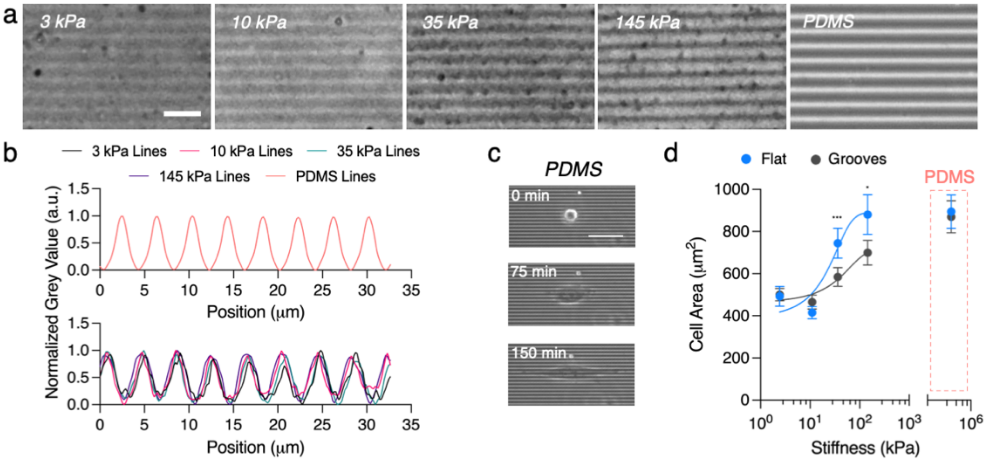


**Fig. S1.** (**A**) Phase contrast image of 2 µm wide and 1 µm high grooves on PAA gels of different stiffness and PDMS. Scale bar, 10 µm. (**B**) Intensity profiles of the microstructured grooves for PDMS and PAA gels. (**C**) Snapshots of NIH 3T3 fibroblasts adhering, elongating and aligning on PDMS grooves. Scale bar, 35 µm. (**D**) Cell area versus substrate stiffness. See Table S1 for the number of cells and experiments. Data points (Mean ± CI) were fitted as an eye-guide. Statistical significance was assessed by Tukey's tests.

**Fig. S2.**

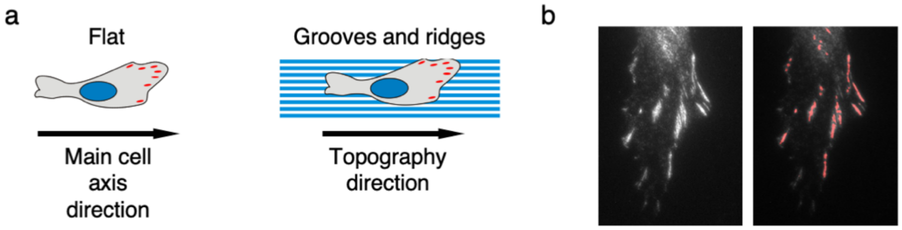


**Fig. S2.** (**A**) Schematics depicting the reference used to evaluate focal adhesion alignment on flat and grooved substrates. (**B**) Representative image of the detection of focal adhesion used to determine area and orientation.

**Fig. S3.**

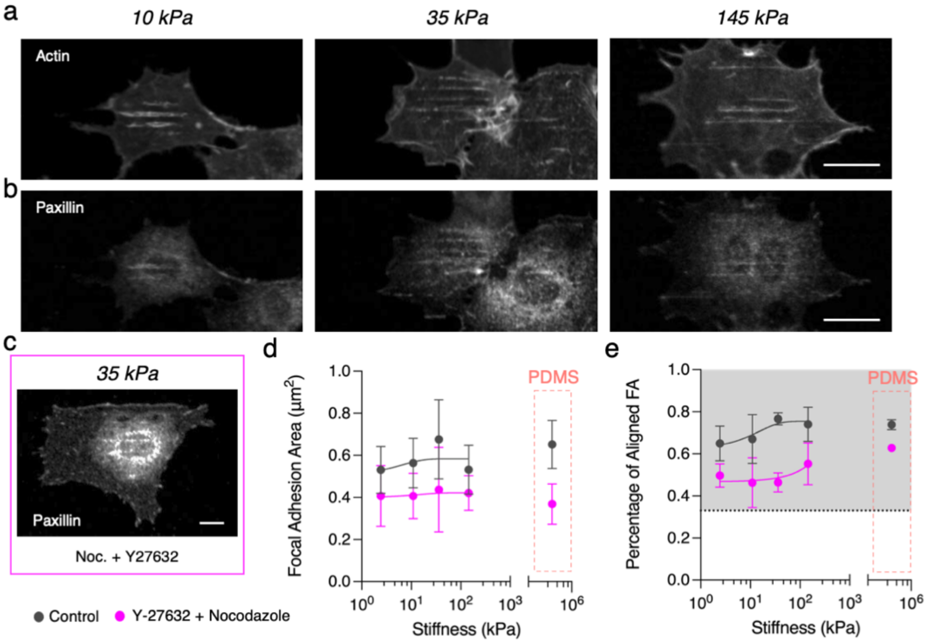


**Fig. S3.** Representative images of cells treated with Y-27632 and nocodazole on grooves of increasing stiffness. (**A**) F-actin (phalloidin) and (**B**) paxillin. Scale bar, 20 µm. (**C**) Paxillin immunostaining of fibroblast treated with 20 µM Y-27632 and 1 µM nocodazole on 2 µm wide grooves of 35 kPa. Scale bar, 20 µm. (**D**) Focal adhesion area and (**E**) percentage of aligned focal adhesions as a function of increasing grooves stiffness. Grey area in (**E**) corresponds to values above the expected for a random distribution. See Tables S9 for the number of cells and experiments.

**Fig. S4.**

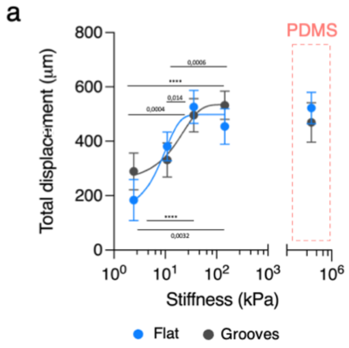


**Fig. S4.** (**A**) Total displacement of cells migrating for 6h on substrates of different stiffness. See Tables S10 for the number of cells and experiments. Data points (Mean ± CI) were fitted as an eye-guide. Statistical significance was assessed by Tukey's tests (**A**).

**Fig. S5.**

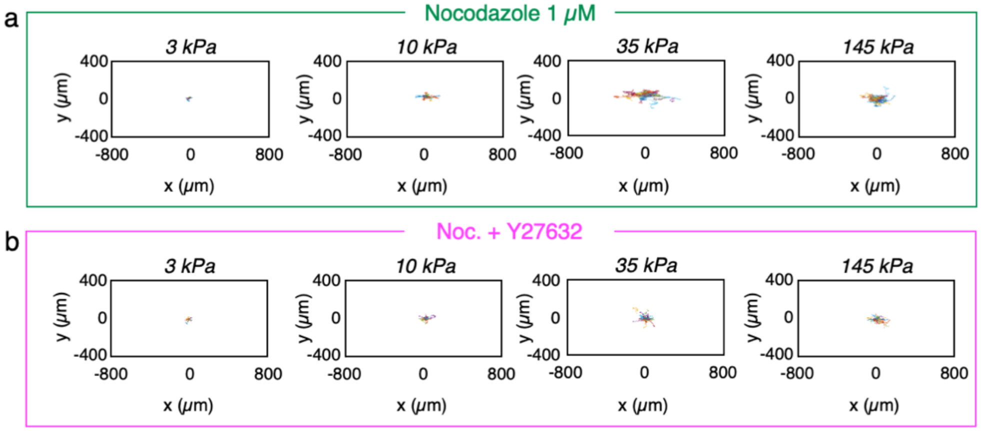


**Fig. S5.** (**A**) Trajectories of 3T3 fibroblasts treated with 1 μM nocodazole and (**B**) 20 μM Y-27632 and 1 μM nocodazole on grooves of increasing stiffness.

**Fig. S6.**

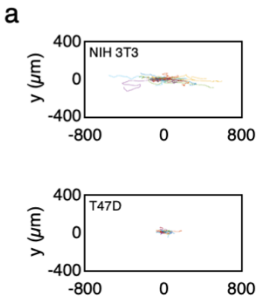


**Fig. S6.** (**A**) Trajectories of NIH 3T3 fibroblasts and T47D single cells migrating on grooved PDMS substrates.

**Fig. S7.**

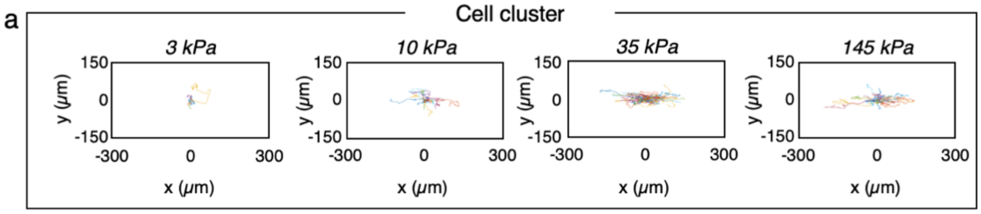


**Fig. S7.** (**A**) T47D clusters' trajectories on grooves of increasing stiffness.

### Table S1.

**Table S1 – NIH 3T3 morphology**

| Stiffness (kPa) | Flat | | Grooves | |
|---|---|---|---|---|
| | N experiments | n cells | N experiments | n cells |
| 3 | 3 | 52 | 5 | 116 |
| 10 | 5 | 57 | 10 | 114 |
| 35 | 6 | 94 | 8 | 134 |
| 145 | 4 | 67 | 5 | 124 |
| PDMS | 4 | 133 | 6 | 184 |

**Table S2.**

**Table S2 – T47D morphology**

| Stiffness (kPa) | Flat | | Grooves | |
|---|---|---|---|---|
| | N experiments | n cells | N experiments | n cells |
| 3 | 2 | 10 | 3 | 37 |
| 10 | 2 | 12 | 4 | 21 |
| 35 | 3 | 17 | 3 | 42 |
| 145 | 2 | 9 | 3 | 45 |
| PDMS | 3 | 42 | 3 | 91 |

### Table S3.

**Table S3 – NIH 3T3 focal adhesions**

| Stiffness (kPa) | Control | | Nocodazole+Y27632 | |
|---|---|---|---|---|
| | N experiments | n cells | N experiments | n cells |
| 3 | 4 | 21 | 2 | 5 |
| 10 | 3 | 24 | 2 | 11 |
| 35 | 4 | 15 | 2 | 4 |
| 145 | 4 | 25 | 2 | 25 |
| PDMS | 5 | 25 | 2 | 11 |

### Table S4.

**Table S4 – NIH 3T3 – Y27632 morphology**

| Stiffness (kPa) | Flat | | Grooves | |
|---|---|---|---|---|
| | N experiments | n cells | N experiments | n cells |
| 3 | 3 | 17 | 3 | 57 |
| 10 | 3 | 14 | 3 | 63 |
| 35 | 4 | 40 | 4 | 79 |
| 145 | 4 | 38 | 4 | 83 |
| PDMS | 3 | 75 | 3 | 196 |

### Table S5.

**Table S5 – NIH 3T3 actin profile**

| Stiffness (kPa) | Flat | | Grooves | |
|---|---|---|---|---|
| | N experiments | n cells | N experiments | n cells |
| 3 | -- | -- | 4 | 31 |
| 10 | -- | -- | 3 | 35 |
| 35 | -- | -- | 3 | 39 |
| 145 | -- | -- | 3 | 36 |
| PDMS | -- | -- | 4 | 28 |

### Table S6.

**Table S6 – NIH 3T3 tubulin profile**

| Stiffness (kPa) | Flat | | Grooves | |
|---|---|---|---|---|
| | N experiments | n cells | N experiments | n cells |
| 3 | -- | -- | 3 | 22 |
| 10 | -- | -- | 3 | 36 |
| 35 | -- | -- | 3 | 39 |
| 145 | -- | -- | 3 | 37 |
| PDMS | -- | -- | 3 | 27 |

### Table S7.

**Table S7 – NIH 3T3 – Nocodazole morphology**

| Stiffness (kPa) | Flat | | Grooves | |
|---|---|---|---|---|
| | N experiments | n cells | N experiments | n cells |
| 3 | 3 | 16 | 3 | 88 |
| 10 | 4 | 51 | 5 | 102 |
| 35 | 3 | 30 | 5 | 95 |
| 145 | 3 | 13 | 4 | 63 |
| PDMS | 3 | 37 | 3 | 137 |

**Table S8.**

**Table S8 – NIH 3T3 – Nocodazole+Y27632 morphology**

| Stiffness (kPa) | Flat | | Grooves | |
|---|---|---|---|---|
| | N experiments | n cells | N experiments | n cells |
| 3 | 3 | 22 | 3 | 47 |
| 10 | 3 | 15 | 3 | 63 |
| 35 | 3 | 28 | 3 | 85 |
| 145 | 3 | 28 | 3 | 84 |
| PDMS | 3 | 39 | 3 | 54 |

### Table S9.

**Table S9 – NIH 3T3 focal adhesions**

| Stiffness (kPa) | Control | | Nocodazole+Y27632 | |
|---|---|---|---|---|
| | N experiments | n cells | N experiments | n cells |
| 3 | 4 | 21 | 2 | 5 |
| 10 | 3 | 24 | 2 | 11 |
| 35 | 4 | 15 | 2 | 4 |
| 145 | 4 | 25 | 2 | 25 |
| PDMS | 5 | 25 | 2 | 11 |

### Table S10.

**Table S10 – NIH 3T3 migration**

| Stiffness (kPa) | Flat | | Grooves | |
|---|---|---|---|---|
| | N experiments | n cells | N experiments | n cells |
| 3 | 4 | 13 | 3 | 24 |
| 10 | 4 | 23 | 5 | 24 |
| 35 | 5 | 31 | 5 | 65 |
| 145 | 4 | 21 | 3 | 69 |
| PDMS | 4 | 30 | 3 | 31 |

### Table S11.

**Table S11 – NIH 3T3 – Nocodazole migration**

| Stiffness (kPa) | Flat | | Grooves | |
|---|---|---|---|---|
| | N experiments | n cells | N experiments | n cells |
| 3 | 4 | 11 | 3 | 15 |
| 10 | 4 | 17 | 3 | 28 |
| 35 | 4 | 28 | 3 | 49 |
| 145 | 3 | 24 | 3 | 38 |
| PDMS | 4 | 25 | 3 | 25 |

### Table S12.

**Table S12 – NIH 3T3 – Nocodazole+Y27632 migration**

| Stiffness (kPa) | Flat | | Grooves | |
|---|---|---|---|---|
| | N experiments | n cells | N experiments | n cells |
| 3 | 2 | 13 | 2 | 32 |
| 10 | 2 | 10 | 2 | 52 |
| 35 | 2 | 26 | 2 | 67 |
| 145 | 2 | 25 | 2 | 46 |
| PDMS | 2 | 16 | 2 | 29 |

### Table S13.

**Table S13 – T47D migration**

| Stiffness (kPa) | Flat | | | Grooves | | |
|---|---|---|---|---|---|---|
| | N experiments | n cells | n cluster | N experiments | n cells | n cluster |
| 3 | 2 | 8 | 19 | 3 | 14 | 7 |
| 10 | 2 | 13 | 16 | 4 | 41 | 19 |
| 35 | 3 | 4 | 5 | 3 | 47 | 66 |
| 145 | 2 | 5 | 13 | 3 | 34 | 34 |
| PDMS | 3 | 18 | 22 | 3 | 45 | 24 |

### Table S14.

**Table S14 – PAA gels formulation**

| Stiffness (kPa) | % Acrylamide | % Bis-acrylamide |
|---|---|---|
| 3 | 7.5 | 0.050 |
| 10 | 7.5 | 0.075 |
| 35 | 12 | 0.150 |
| 145 | 12 | 0.600 |

**Movie S1.**

NIH 3T3 fibroblasts spreading on topographical grooves and ridges made of PAA gels of different stiffness (3 – 145 kPa). Time hh:mm. Scale bar 50 µm.

**Movie S2.**

NIH 3T3 fibroblasts migrating on topographical grooves and ridges made of PAA gels of different stiffness (3 – 145 kPa). Time hh:mm. Scale bar 50 µm.

**Movie S3.**

T47D single cell migrating on 35kPa topographical grooves and ridges. Time hh:mm. Scale bar 50 µm.

**Movie S4.**

T47D cell cluster migrating on 35kPa topographical grooves and ridges. Time hh:mm. Scale bar 50 µm.
